# Intrinsically Disordered Regions Form Nucleoli and Cajal Bodies While Fostering RNA Modification

**DOI:** 10.1101/2025.06.18.660405

**Authors:** Koceila Meznad, Manisha Deogharia, Ludivine Wacheul, Christiane Zorbas, Denis L.J. Lafontaine, U. Thomas Meier

## Abstract

One of the densest compartments in the cell is the dense fibrillar component (DFC) of the nucleolus, consisting mainly of nascent ribosomal RNA (rRNA), small nucleolar ribonucleoproteins (snoRNPs) and their chaperone Nopp140. How this biomolecular condensate is formed and what underlies its structure is poorly understood like that of most liquid-liquid phase separated condensates. Although we established that Nopp140 is important for the cohesiveness of the DFC and for rRNA modification, it is not known how this is achieved. Here we demonstrate that Nopp140 concentrates intrinsically disordered and nuclear localization signal (NLS)-rich protein regions (IDRs), including a newly identified RNA polymerase I C-terminal domain (CTD) of the RNA polymerase I associated factor PAF49. Altogether, this network forms the DFC, a liquid-liquid phase separated biomolecular condensate that fosters rRNA modification. This mechanism ensures near 100 percent modification efficiency at some 200 nucleotides in every one of the 10 million or so rRNAs per cell.

## INTRODUCTION

Ribosome synthesis is initiated by transcription of ribosomal DNA by RNA polymerase I (pol I). Trains of ∼100 molecules transcribe each active rDNA gene generating an equal number of growing nascent pre-rRNAs, famously depicted in electron micrographs as “Christmas trees” by Oscar Miller (Miller and Beatty, 1969). The fragments of each transcript that are destined for incorporation into mature ribosomes are site-specifically modified at some 200 nucleotides. A similar number of H/ACA and C/D small nucleolar ribonucleoproteins (snoRNPs) insert pseudouridines and 2’-O-methyl groups, respectively (Kiss, 2001; Decatur and Fournier, 2002; Meier, 2005; Watkins and Bohnsack, 2012). Together with ribosome assembly factors and Nopp140, this enormous mix of proteins and RNAs forms the dense fibrillar component (DFC) of the nucleolus. Although the individual components are well established, how the DFC coalesces to form a biomolecular condensate is unknown. We now describe in molecular detail how these proteins, RNAs, and ribonucleoproteins form a phase conducive to rRNA modification without missing a site (Meier, 2022).

By phase contrast microscopy, the nucleolus is the densest part of a mammalian cell clearly distinct from the surrounding nucleoplasm. Transmission electron microscopy reveals three nucleolar subcompartments, multiple lighter staining fibrillar centers (FCs) containing rDNA genes and pol I molecules not engaged in transcription (Thiry et al., 2000). FCs are surrounded by the darkest staining dense fibrillar component (DFC). Transcription occurs at the interphase between FC and DFC with the nascent pre-rRNAs extending into the DFC (Lafontaine et al., 2021; Dundr and Misteli, 2001). After transcription termination and pre-rRNA cleavage, pre-ribosomes are released into the granular component (GC) of the nucleolus for further assembly and folding before transport through the nucleoplasm across nuclear pore complexes (NPCs) into the cytoplasm to function in mRNA translation (Hernandez-Verdun et al., 2010; Scheer and Hock, 1999).

Ribosomal RNA is modified in the DFC where all snoRNPs are housed. Modification must occur cotranscriptionally because many of the modified nucleotides are buried inside the folded rRNA inaccessible to the bulky snoRNPs. Additionally, some modifications are present in double stranded regions of folded rRNA preventing hybridization access to guide snoRNAs after folding. As rRNA is modified at most sites to near full extent (Birkedal et al., 2015; Taoka et al., 2018), the process must be enormously efficient for all snoRNPs to scan the entire 13kb of pre-rRNA for modification sites. This could be achieved by direct recruitment to the elongating polymerase or by high concentration of snoRNPs around nascent pre-rRNA or a combination thereof. Either way will require crowding of snoRNPs resulting in the high density of the DFC. SnoRNPs come in two flavors based on their guide RNAs, H/ACA and C/D. Each RNP consists of a duplex of four core proteins and one of hundreds of snoRNAs. H/ACA RNPs harbor the pseudouridine synthase NAP57 (aka dyskerin or Cbf5), that is stabilized by the short NOP10 protein forming a landing pad for NHP2; GAR1 associates independently with NAP57. C/D RNPs encompass the methyltransferase fibrillarin, the two related NOP56 and NOP58 core proteins, together with the 15.5K protein that is related to NHP2 of H/ACA RNPs. The RNPs are highly charged with basic proteins and the acidic RNA. Therefore, it requires a charged protein to neutralize their electrostatic attraction and repulsion.

Nopp140 a protein of the DFC and Cajal bodies fulfills this requirement. It has ten alternating acidic serine patches separated by exclusively basic stretches. All serines in those patches are phosphorylated by casein kinase 2 (CK2) causing a shift in migration of 40kD on an SDS-PAGE (Meier and Blobel, 1992). Although we know for many years that Nopp140 can associate with both types of snoRNPs in a phosphorylation dependent manner (Wang et al., 2002), it is not clear how that is possible given the two disparate types of snoRNPs. We now show the importance of intrinsically disordered regions (IDRs) of NAP57, NOP56, and NOP58, and of the newly defined C-terminal domain of the pol I subunit PAF49 for interaction with Nopp140.

Additionally, glycine-arginine-rich (GAR) domains of GAR1 and fibrillarin contribute to interaction with each other and with Nopp140. Finally, the transient and short-lived base pairing between snoRNAs and rRNAs contribute to a liquid-liquid phase separated body that forms the DFC. In case of Cajal bodies (CBs), the CB marker protein and Nopp140 interactor coilin replaces pol I while spliceosomal small nuclear RNAs (snRNAs) replace rRNA. Meanwhile, small CB ribonucleoproteins (scaRNPs), nearly identical to snoRNPs, perform the modification of snRNAs.

## RESULTS

### Nopp140 binds to H/ACA and C/D RNPs through the IDRs of their core proteins

Both, H/ACA and C/D RNPs bind to Nopp140 but is not known how (Yang et al., 2000). In fact, the pseudouridine synthase NAP57 of H/ACA RNPs and the C/D core protein NOP58 (aka NAP65) were identified in association with Nopp140 (Yang et al., 2000; Meier and Blobel, 1994). The only commonality between H/ACA and C/D RNPs are the intrinsically disordered C-terminal tails of their core proteins NAP57, NOP58, and NOP56 with similarly spaced SV40 large T antigen-type classic monopartite nuclear localization signals (NLSs) (Fig. 1A). The intrinsic disorderedness of all domains designated as such in this manuscript is supported by several algorithms including AlphaFold (Jones and Cozzetto, 2015; Erdős et al., 2021; Jumper et al., 2021). The NLSs of the IDRs are separated by spacers of 18 ±1.7 (average ± standard deviation), 18 ±5.2, and 21.6 ±2.6 dotted with negative charges for NAP57 (3; number of NLSs in parentheses), NOP58 (4), and NOP56 (6), respectively. NAP57 has an additional short N-terminal IDR with a single NLS. Since we identified Nopp140 as an NLS-binding protein (Meier and Blobel, 1990), we tested for its interaction with the NLS containing tails using amylose resin pulldowns of recombinant proteins tagged with maltose binding protein (MBP; Fig. 1). Nopp140 and its derived constructs were tagged with a hexa histidine tag for nickel affinity purification and expressed from baculovirus in insect cells to be fully phosphorylated. MBP-NAP57 was pulled down by with amylose resin (Fig. 1B, lane 4) and bound Nopp140 (lane 6), which did not sediment alone (lane 5). Proteins were treated with micrococcal nuclease or RNase A indicating that nucleic acid was not involved in NAP57-Nopp140 interaction. Note, MBP-NAP57 pulldown of Nopp140 was included as a reference in many of the gels throughout this study. The intrinsically disordered region (IDR) of NAP57 was necessary for Nopp140 binding, as only full-length NAP57 (Fig. 1C, lane 7) and constructs with its C-terminal or N-terminal NLS IDRs (lanes 8 and 9, respectively) pulled down Nopp140 but not NAP57 without both tails (lane 10) or a control protein, MS2 coat protein (MCP, lane 11). Nopp140 alone did not bind to amylose beads (Fig. 1C, lane 12). On the other hand, the C-terminal NLS IDR alone was sufficient for Nopp140 pulldown (Fig. 1D, lane 6) but not its short N-terminal IDR alone (lane 5). Additionally, the C-terminal NLS IDRs of NOP58 and NOP56 bound to Nopp140 (Fig. 1E, lanes 5 and 6). Again, the NLS IDR of NOP56 was necessary (Fig. 1F, lane 4) and sufficient (Fig. 1G, lane 4) for Nopp140 binding. Thus, Nopp140 can interact with the NLS containing IDRs of core proteins of both, H/ACA and C/D RNPs, providing a molecular basis for its physical association with both RNPs.

**Figure 1.**
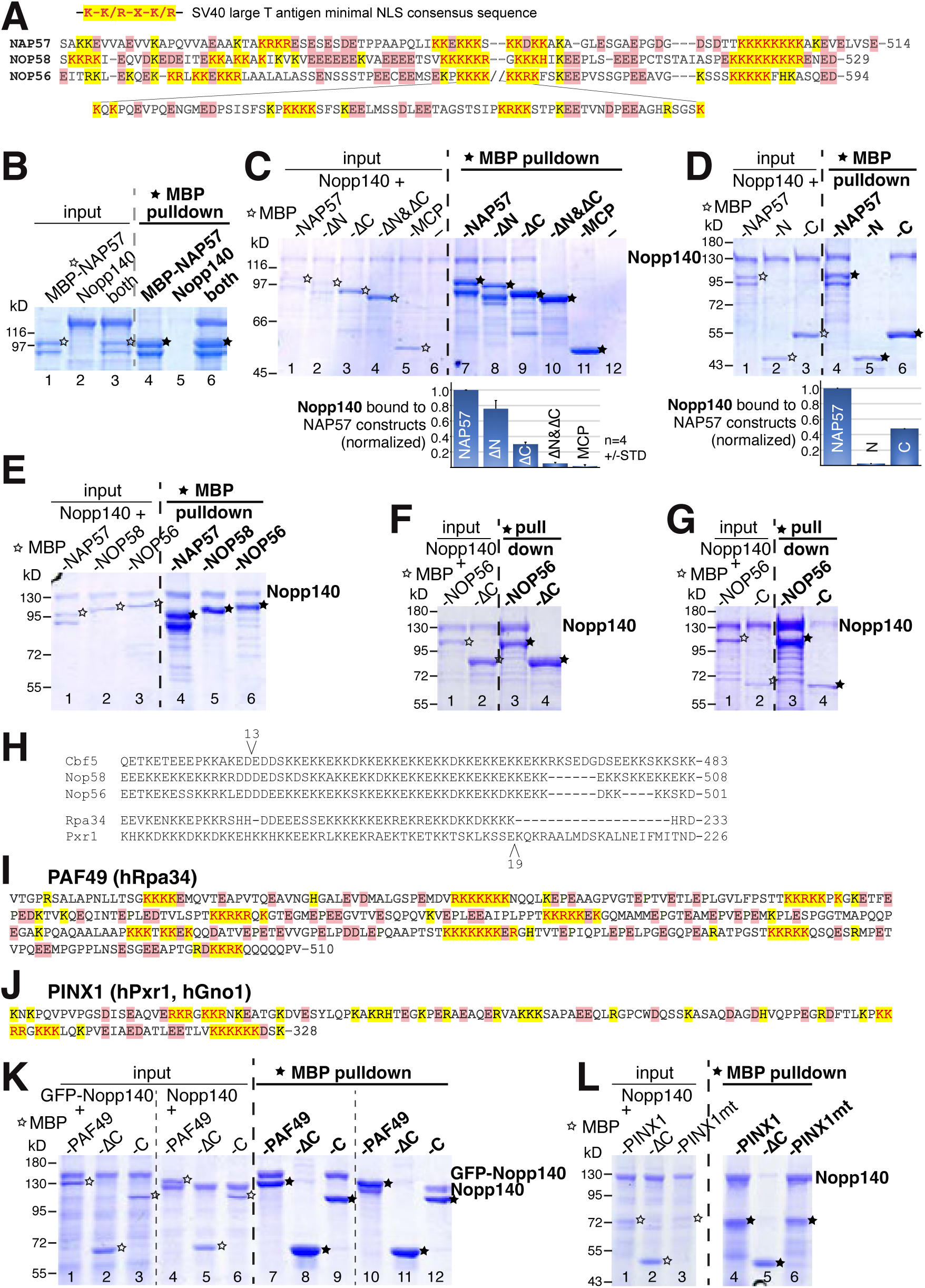
Charged, nuclear localization signal-rich, intrinsically disordered regions of snoRNP core proteins and of the RNA polymerase I associated factor PAF49 bind to Nopp140. (A) Amino acid sequences of the C-terminal IDRs of NAP57, NOP58, and NOP56. The minimal SV40 Tag NLS is highlighted above (yellow with red lettering) and indicated in the IDRs. Negatively charged amino acids are highlighted red. (B) Pulldown experiments with amylose beads that bind maltose binding protein (MBP) and analyzed on coomassie blue stained polyacrylamide gels. The left-hand side always depicts the input fraction before pulldown (right-hand side). MBP-NAP57 was expressed in bacteria (lane 1) and phosphorylated Nopp140 in baculovirus (lane 2); both together (lane 3). Nopp140 alone is not pulled down (lane 5), only in association with MBP-NAP57 (lane 6). (C) Nopp140 is pulled down by full-length MBP-NAP57 (lane 7), but less without its N- or C-terminal NLS IDR (lane 8 and 9, respectively). However, MBP-NAP57 without both IDRs fails to pull down Nopp140 (lane 10). Similarly, MBP fused to the viral MS2 coat protein (MCP) fails to pull down Nopp140 (lane 11) and neither is Nopp140 alone (lane 12). Below, the pulldowns are quantified and normalized to MBP-NAP57 including standard deviations from 4 experiments (lanes 7-12). (D) The C-terminal IDR of NAP57 (92aa) pulls down Nopp140 (lane 6) at about 50% of full-length NAP57 (lane 4, quantified below). Whereas the N-terminal IDR (30aa) with only a single NLS barely pulls down Nopp140 (lane 5, quantified underneath). (E) MBP fusions of the C/D core proteins NOP58 (lane 5) and NOP56 (lane 6) pulldown Nopp140 like the H/ACA core protein NAP57 (lane 4). (F) Only full-length MBP-NOP56 (lane 3) but not without its C-terminal IDR (lane 4) pulls down Nopp140. (G) The C-terminal IDR of NOP56 is sufficient for Nopp140 pulldown. (H) Identification of yeast genes with homologous charged IDRs. In blast searches, we used the C-terminal IDR of Cbf5, the yeast ortholog of human NAP57/dyskerin, to identify the C-terminal IDRs of Nop58 and Nop56, the NOP58 and NOP56 orthologs, and of Rpa34 and Pxr1, the only yeast proteins with mammalian homologs. The numbers above and below the aligned sequences signify the location and number of amino acid inserts. (I) Sequence of the 350aa-long charged C-terminal IDR with 9 NLSs of the human Rpa34 homolog PAF49. Same color scheme as in (A). (J) The 138aa-long C-terminal IDR with 3 NLSs of the human Pxr1 homolog PINX1. (K) MBP pulldowns with full-length PAF49 (lanes 7 and 10), PAF49 without its NLS-IDR (lanes 8 and 11), and the PAF49 NLS-IDR alone (lanes 9 and 12). Nopp140 (lanes 10-12) and GFP-Nopp140 (lanes 7-9) were used to better distinguish migration of the protein bands. Lanes 1-6 are inputs. (L) Only full-length PINX1 pulls down Nopp140 (lane 4), but not without its NLS-IDR (lane 5). Mutation of the G patch motif does not impact Nopp140 binding (lane 6).

### Identification of an RNA polymerase I CTD in its associated factor PAF49

To search for additional proteins with charged NLSs IDRs, we used a phylogenetic detour through yeast. The IDRs of the yeast orthologs of NAP57, NOP58, and NOP56, i.e., Cbf5, Nop58, and Nop56, respectively, are conserved on an amino acid sequence level with KKE/D repeats (Fig. 1H). Indeed, a blast search using the C-terminal IDR of Cbf5 yielded two additional proteins with mammalian homologs Rpa34 and Pxr1 (Fig. 1H). Rpa34 corresponds to the human RNA polymerase I associated factor PAF49 with a 350 amino acid long C-terminus containing 9 NLSs separated by stretches dotted with negatively charged amino acids (Fig. 1I). Pxr1 corresponds to the nucleolar G patch helicase cofactor PINX1 with a 140 amino acid long IDR containing 3 NLSs interspersed with negative charges (Fig. 1J). Full-length PAF49 efficiently pulls down GFP-Nopp140 and Nopp140 alone (Fig. 1K, lanes 7 and 10), whereas removal of its NLS IDR abolishes this interaction (lanes 8 and 11) and the IDR alone was sufficient for Nopp140 binding (lanes 9 and 12). Similarly, PINX1 pulls down Nopp140 (Fig. 1L, lane 4) but not without its IDR (lane 5). Mutations of its G patch motif that abolish interaction with the Prp43p/DHX15 helicase (Chen et al., 2014) have no impact on its Nopp140 association (Fig. 1L, lane 6). In summary, the IDRs with NLSs separated by negatively charged spacers are necessary and sufficient for Nopp140 interaction. What domain(s) of Nopp140 bind to these charged NLS IDRs?

### Nopp140 binds to NLS-containing IDRs with its phosphorylated repeat domain

Human Nopp140 is a 699 amino acid long protein that contains 10 central acidic serine repeats of 15.7 ±1.6 amino acids (avg ±std) separated by lysine, alanine, and proline-rich stretches of 32.3 ±7.2 residues (Fig. 2A). All serines in the repeats are phosphorylated by casein kinase 2 (Meier and Blobel, 1992; Wang et al., 2002) and presented the most likely target for NLS interaction. To obtain phosphorylated Nopp140, its cDNA was expressed in Sf9 insect cells with the help of baculovirus and purified via a hexa histidine tag (Fig. 2B, lane 2). Phosphorylation was confirmed by a downshift in migration on SDS-PAGE by some 40kD after treatment with calf intestinal alkaline phosphatase (CIP; Fig. 2B, lane 3), as observed previously (Meier and Blobel, 1992). When mixed with MBP-NAP57 (lanes 4 and 5), only phosphorylated (lane 6) but not (or barely) dephosphorylated Nopp140 (lane 7) was pulled down with amylose beads. This suggested that NAP57 interacted with the phosphorylated and intrinsically disordered repeat domain of Nopp140. Indeed, phosphorylated repeats 1-10 alone were pulled down by MBP-NAP57, -NOP56, and -NOP58, but not MBP alone (Fig. 2C, lanes 7-10). To test how many Nopp140 repeats were necessary for interaction, we tested individual repeats by truncation. Even 3 repeats were pulled down to various degrees by MBP-PAF49 (Fig. 2D) and -NOP58 (Fig. 2E). Also, repeats 8-10 and repeat 1 alone were sufficient for interaction with MBP-NOP58 (Fig. 2F). Apparently, each repeat can bind to an NLS IDR. As with full length Nopp140, interaction of the truncated repeats was dependent on phosphorylation, because dephosphorylation with CIP abolished pulldown by MBP-PAF49 (Fig. 2G, lanes 8, 10, and 12). To assure that dephosphorylation of Nopp140 on its own did not cause aggregation and sedimentation in our assays, we tested pulldowns minus/plus CIP with MBP-MCP (Fig. 2H, lanes 7 and 8) in addition to MBP-NAP57 (lanes 5 and 6). Moreover, we abolished all casein kinase 2 consensus sites in Nopp140 by substituting 46 serines with alanines generating a NoppSA mutant that migrates at 110kD almost with the same mobility as dephosphorylated Nopp140 (Fig. 2I, lanes 1-3). Surprisingly, after CIP treatment, NoppSA migrated even faster suggesting additional phosphorylation sites in Nopp140 besides casein kinase 2 (Fig. 2I, lane 4). Dephosphorylation of Nopp140 mostly abolished interaction with MBP-NAP57 (Fig. 2I, lane 6). Phosphorylation dependence was further supported by the lack of NoppSA pulldown (Fig. 2I, lane 7), which was not altered after dephosphorylation of NoppSA by CIP (lane 8). In conclusion, each of the 10 Nopp140 repeats can interact with an NLS IDR in a phosphorylation dependent manner.

**Figure 2.**
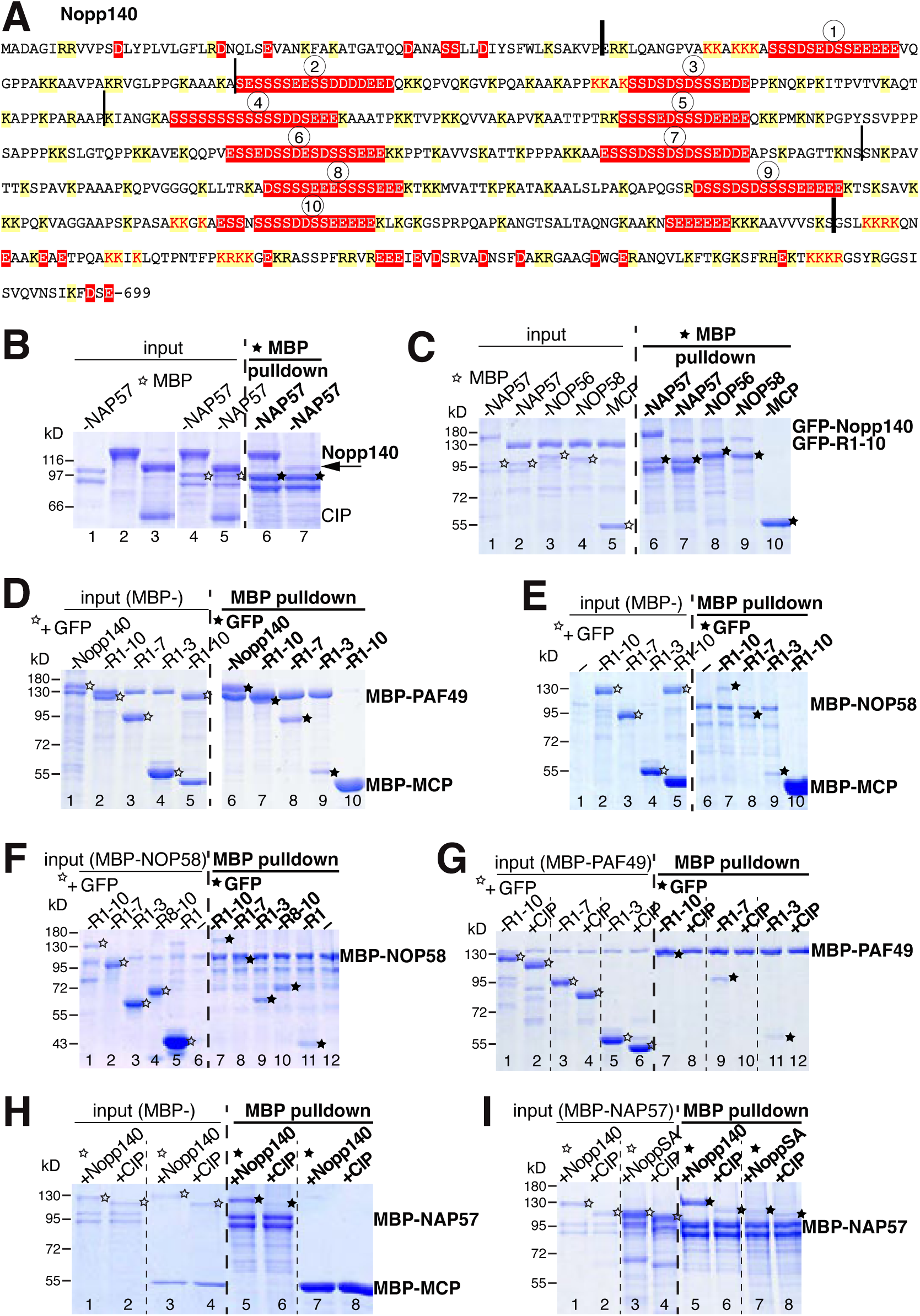
Dissection of Nopp140 requirements for interaction. (A) Amino acid sequence of human Nopp140. Negatively (red) and positively (yellow) charged amino acids are shaded. The 10 acidic serine stretches are numbered. All serines in those stretches are or become casein kinase 2 consensus sites after phosphorylation. Consensus NLSs are indicated (red lettering). The N- and C-termini are demarcated from the large central repeat domain (thick black bars). The borders of the repeats tested in the figure are marked (thin black lines). All Nopp140 constructs were expressed fully phosphorylated in baculovirus-transduced Sf9 insect cells. All others were expressed in bacteria. (B) MBP-NAP57 (lane 1), Nopp140 (lane 2), calf intestinal phosphatase (CIP, lower band) added to Nopp140 (lane 3). Note the near 40kD shift in migration upon dephosphorylation of Nopp140 (lane 2 vs. 3). MBP-NAP57 added to Nopp140 (lanes 4 and 5). Only phosphorylated Nopp140 (lane 6) but barely dephosphorylated Nopp140 (lane 7) is pulled down by NAP57. (C) MBP-NAP57 and C/D core proteins -NOP56 and -NOP58 pull down the Nopp140 repeat domain alone (lanes 7-9), but not the control MBP-MCP (lane 10). For identification purposes on SDS-PAGE, GFP-Nopp140 fusion constructs were used instead of Nopp140 alone. (D) MBP-PAF49 also pulled down full-length Nopp140 (lane 6), its repeat domain (R1-10, lane 7), or repeats 1-7 (lane 8), or 1-3 (lane 9). (E) Similarly, MBP-NOP58 pulled down repeat domains 1-10 (lane 7), 1-7 (lane 8), and 1-3 (lane 9). (F) MBP-NOP56 pulled down the same repeats of Nopp140 (lanes 7-9) and was additionally tested for repeats 8-10 (lane 10) and repeat 1 alone (lane 11), which were all pulled down. (G) All tested repeats were only pulled down in a phosphorylation dependent manner. MBP-PAF49 pulled down repeats 1-10 (lane 7), 1-7 (lane 9), and 1-3 (lane 11), but not after dephosphorylation (lanes 8, 10, and 12). (H) Nopp140 alone was not pulled down by MBP-NAP57 after dephosphorylation (lane 6), and MBP-MCP did not pull down Nopp140, whether phosphorylated (lane 7) or not (lane 8). (I) Nopp140 before and after CIP treatment (lanes 1 and 2, respectively) and Nopp140 with 46 CK2 consensus site serines replaced by alanines in the repeat domain (NoppSA, lane 3). Surprisingly CIP treatment of NoppSA further accelerated its mobility (lane 4) indicating additional phosphorylation sites. Regardless, whereas Nopp140 was efficiently pulled down by MBP-NAP57 (lane 5), CIP treated constructs were not or only barely pulled down (lanes 6-8).

### Nopp140 mediates interaction between the NLS IDR of PAF49 and those of other proteins

To test if Nopp140 could bridge two proteins with different NLS IDRs, we added MBP-PAF49 and NOP56 to 3 samples (Fig. 3A, lanes 1-3) and supplemented one sample with GFP-Nopp140 (lane 2, star) and one additionally with CIP to dephosphorylate Nopp140 (lane 3, star). Whereas MBP-PAF49 failed to pulldown NOP56 alone (lane 4), addition of GFP-Nopp140 stimulated its pulldown (lane 5, stars) whereas CIP treatment abolished all interactions (lane 6). In a reverse manner, we added MBP-NOP58 to 3 samples (Fig. 3B, lanes 1-3) and PAF49 to two samples (lanes 2 and 3). Nopp140 was added to sample 1 and 3 (Fig. 3B, lanes 1 and 3, stars). As seen above (Fig. 1E, lane 5), MBP-NOP58 efficiently bound Nopp140 (Fig. 3B, lane 4, star) but barely PAF49 alone (lane 5). However, addition of Nopp140 significantly increased pulldown of PAF49 (lane 6, stars) demonstrating a ternary complex between Nopp140, PAF49, and NOP58. These findings supported our model whereby the CTD of pol I, i.e., the NLS IDR of PAF49, recruits Nopp140 and with it both families of snoRNPs through the NLS IDRs of their core proteins.

**Figure 3.**
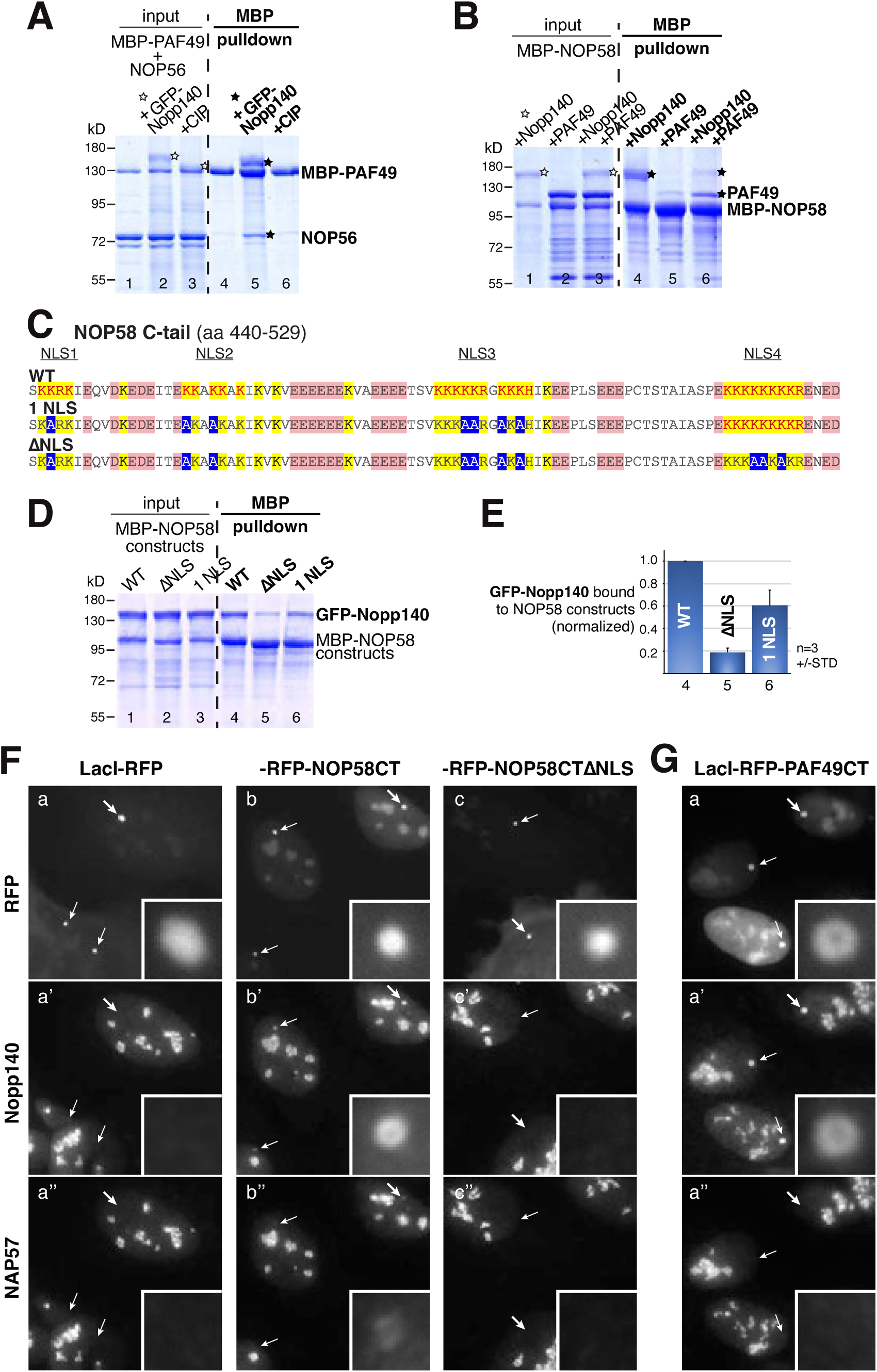
Nopp140 interacts with NLS-IDRs in vitro and in vivo. (A) Nopp140 forms a ternary complex between PAF49 and NOP56. All samples contain MBP-PAF49 and NOP56 (lanes 1-3), plus GFP-Nopp140 (lanes 2 and 3) and CIP (lane 3). MBP-PAF49 alone does not pull down NOP56 (lane 4), only in the presence of Nopp140 (lane 5) but not after dephosphorylation (lane 6). (B) MBP-NOP58 pulls down Nopp140 (lane 4), but barely PAF49 (lane 5) which is significantly increased in the presence of Nopp140 (lane 6) in a ternary complex. (C) Wild type (WT) NLS-IDR of NOP58 contains 4 NLSs (red lettering). Strategic replacement of 7 lysines by alanines (blue) leaves one NLS (1 NLS). Substitution of 10 lysines abolishes all NLSs (ΔNLS). (D) Relative to WT MBP-NOP58 (lane 4), the ΔNLS MBP-NOP58 construct only pulls down about 20% of GFP-Nopp140 (lane 5) whereas 1NLS was sufficient to pull down more than half of GFP-Nopp140 (lane 6). (E) The amount of GFP-Nopp140 pulled down and quantified from three experiments (average ± standard deviation). (F) Nuclear tethering assay of Lac repressor bound to Lac operator arrays inserted at one location of the genome of U2OS cells and transiently transfected with a LacI-RFP fusion construct (a). Indirect immunofluorescence with Nopp140 (a’) and NAP57 (a’’) constructs (panel width 51µm). Integration sites are marked by arrows; the thicker arrow is 20x enlarged in the inset (2.6µm). (b-b’’) the NOP58 C-terminus was fused to LacI-RFP and (c-c’’) the NOP58 C-terminus ΔNLS was fused to LacI-RFP. Note the strong binding of Nopp140 to the integration site (b’) and even a slight binding by NAP57 (b’’), but no binding of Nopp140 to the ΔNLS constructs. (G) Fusion of the PAF49 C-terminus to LacI-RFP (a) also recruits Nopp140 (a’) but not NAP57 (a’’).

We employed the NLS-IDR of NOP58 to test whether it was indeed the NLSs of the IDR that interacted with the NLS-binding protein Nopp140. There are four monopartite NLSs of the minimal -K-K/R-X-K/R-SV40 large T antigen-type in the IDR of NOP58, with NLS 2 to 4 harboring at least duplicate NLSs (Fig. 3C, WT). To abolish the NLSs, we strategically substituted critical lysines by alanines (Fig. 3C, blue) that constitute the minimal monopartite classical NLS required for nuclear protein import (Kalderon et al., 1984; Lanford and Butel, 1984; Chelsky et al., 1989). Thus, NOP58 was generated with no NLS (Fig. 3C, ΔNLS) or with a single NLS (1 NLS). When expressed in bacteria and used in pulldown experiments of Nopp140, the NOP58ΔNLS interacted 5-fold less with Nopp140 (Fig. 3D, lanes 4 and 5), whereas NOP58 with a single NLS restored binding to more than half of the wild type (lane 6). Averages of these data from multiple experiments were quantified (Fig. 3E). Consequently, Nopp140 can coalesce multiple snoRNPs and ribosome biogenesis factors at RNA polymerase I via the PAF49 NLS-IDR.

### NLS-dependent interactions with Nopp140 occur in the nucleus

Up to this point, all interactions were demonstrated with recombinant purified proteins. Hence it was important to replicate the findings in the cell nucleus, in the presence of endogenous nuclear proteins. For this purpose, we employed our previously developed nuclear tethering assay, which takes advantage of a U2OS cell line that has an array of Lac operator sequences (LacO) integrated at one site of its genome (Janicki et al., 2004; Darzacq et al., 2006). Upon transfection of the Lac inhibitor fused to the monomeric red fluorescent protein (LacI-RFP), it bound to the LacO arrays yielding a bright red fluorescent signal (Fig. 3F a, arrows) that did not accumulate Nopp140 (a’) nor NAP57 (a”). However, appending the wild type C-terminal NLS-IDR of NOP58 (Fig. 3C), concentrated endogenous Nopp140 (Fig. 3F a’, arrows and inset) and even some NAP57 (a’’), presumably through binding to Nopp140 in a ternary complex. Nopp140 interaction with the NOP58CT was lost if its 4 NLSs were abolished by strategic substitution of 10 lysines by alanines (Fig. 3C and F c’). Nopp140 was similarly recruited to the LacO integration sites if the CT of PAF49 with its 9 NLSs was fused to LacI-RFP (Fig. 1I and 3G a’).

This recruitment of Nopp140, but barely NAP57, to NLS IDRs, may be like Nopp140, but not NAP57, recruitment to pseudoNORs (Prieto and McStay, 2007). Thus, the in vitro established NLS-dependent interaction of Nopp140 with the NLS-IDRs was confirmed in vivo.

### Removal of the PAF49 IDR disrupts the tripartite organization of the nucleolus

To investigate the role of the pol I CTD, i.e., the charged IDR of PAF49 with 9 NLSs, we used CRISPR/Cas9 technology to replace the IDR fully or partially by GFP. This was performed in near haploid HAP1 cells (Carette et al., 2011) with the following versions of PAF49, a) endogenous PAF49, b) full-length PAF49 tagged with GFP, c) PAF49 with its IDR replaced by GFP, and d) PAF49 with half of its IDR replaced by GFP leaving 4 NLSs (Fig. 4A). Analysis of the four cell lines by fluorescence microscopy shows the integration of GFP in a nucleolar pattern identical to untagged PAF49 (Fig. 4B, a and b) and a reduction of PAF49 signal after replacement of all or parts of its IDR (c and d). Apparently, partial or full replacement of the IDR caused a destabilization of PAF49. PAF49 antibodies appear to recognize an epitope in the N-terminal half of the IDR as no signal was observed in the cells without IDR (Fig. 4B, c). Nopp140 distribution in the nucleolus seemed unperturbed at this resolution. These results were confirmed by western blotting of whole cell extracts probed with GFP and PAF49 antibodies (Fig. 4C and D). Moreover, in the stable cell lines, Nopp140 (Fig. 4E) and core components of H/ACA (F) and C/D (G) snoRNPs were not affected by GFP integration into the PAF49 gene. All four cell lines grow and divide at the same rate indicating that ribosome synthesis must not be affected (not shown).

**Figure 4.**
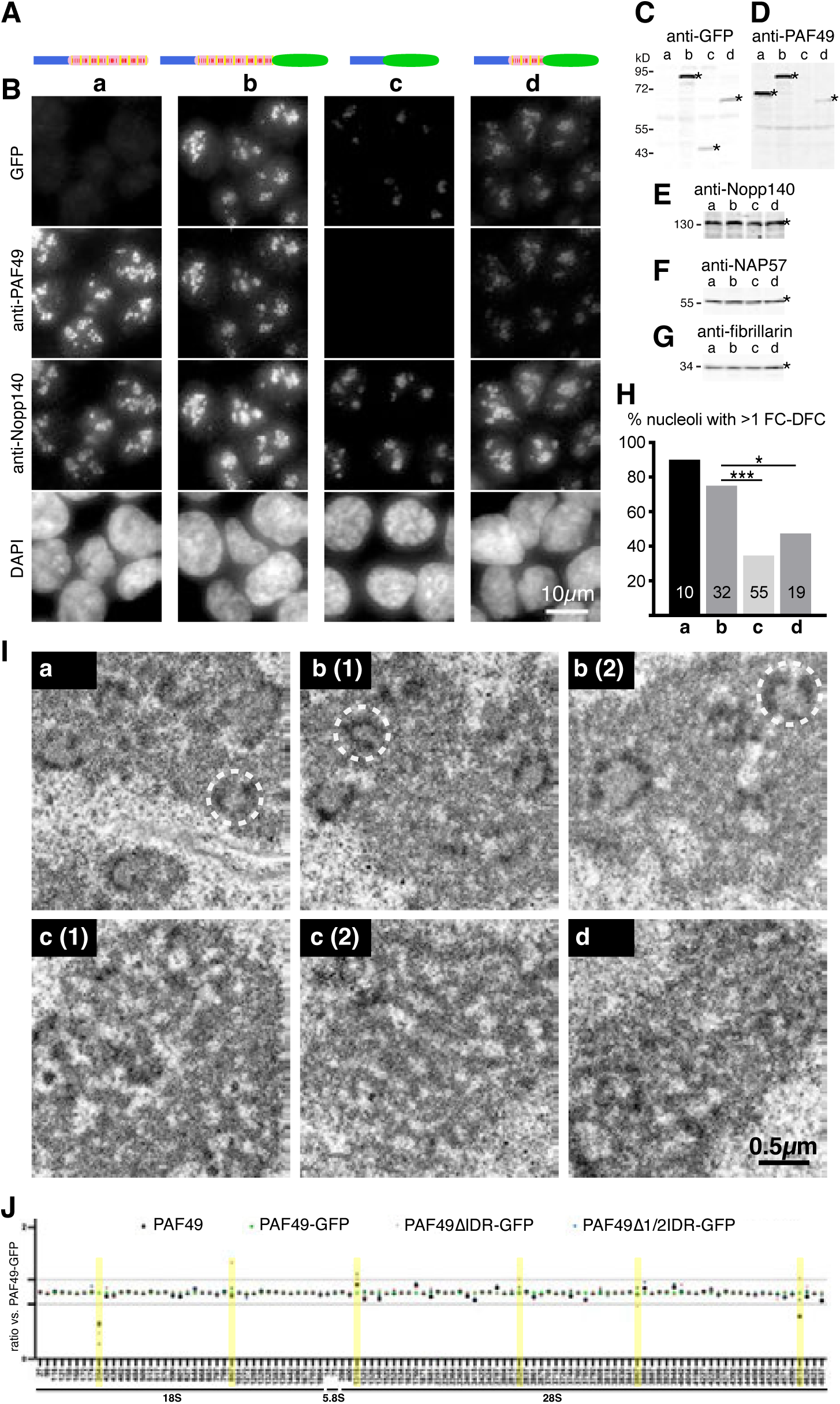
The PAF49 C-terminus is required for nucleolar organization but not rRNA modification. (A) Schematic representation of the PAF49 CRISPR constructs in the HAP1 cells: a) wild type PAF49, b) PAF49 with GFP appended, c) GFP replacing the IDR, and d) GFP partially replacing the PAF49 IDR. The constructs are drawn to scale (blue: structured part of PAF49, pink-yellow-red: IDR). (B) (Immuno) fluorescence of the antigen indicated on the left of the cells indicated in (A). DNA stain (DAPI) was used to mark the nuclei. Note, GFP addition does not change the nucleolar localization of PAF49 (b) but seems to destabilize the constructs without and partial IDR (c and d). The epitope of the PAF49 antibodies must be mostly in the PAF49 IDR because the signal is abolished when it is replaced by GFP (c). (C) The fluorescence data is confirmed on western blots of whole cell extracts probed with GFP (C) and PAF49 (D) antibodies, i.e., no GFP protein band in wild type HAP1 cells (C, a), but after GFP insertion (b), and reduced after IDR replacement (c) or partial replacement (d). (D) GFP insertion does not alter the amount of PAF49 (compare lanes a and b), but PAF49 antibodies only partially recognize the truncated IDR construct (lane d) and not at all after replacement (lane c). Note the bands recognized by the antibodies are marked by asterisks. Remarkably, the amounts of Nopp140 (E), H/ACA RNPs (F, NAP57), and C/D RNPs (G, fibrillarin) do not vary between the cell lines. (I) Transmission electron micrographs of nucleoli from the four cell lines (a-d). Two independent clones of cell lines (b) and (c) are depicted. Note the classic nucleolar tripartite structure with multiple units of fibrillar centers (FCs) surrounded by dense fibrillar components (DFCs) embedded in the granular component (GC) in cells with wild type PAF49 (a) and GFP fused to it (b). However, this organization is disrupted in cells where the PAF49 IDR was partially (d) or wholly (c) replaced by GFP. (H) Counting the FC-DFC units per nucleolar section (one is circled in (I)) shows a significant difference between wild type and IDR deleted or partially deleted cells *** p < 0.001 and * p < 0.05. (J) RiboMethSeq analysis reveals that only 6 out of 107 2’-O-methylation sites in rRNA vary by more than 1.2-fold between the four cell lines. Additionally, the affected sites (yellow) do not show a pattern that coincides with a cell line, e.g., methylation in the IDR deleted cells is upregulated at nucleotide 687 of 18S but down regulated at position 3867 of 28S rRNA.

To study the impact on nucleolar organization at higher resolution, we analyzed all cell lines by transmission electron microscopy (Fig. 4I). In parent HAP1 cells, the nucleolus exhibits the classic tripartite structure of several lightly stained and amorphous fibrillar centers (FCs) surrounded by the darker staining dense fibrillar component (DFC), which are altogether embedded in the granular component (Fig. 4I a, one FC-DFC unit is highlighted by a dashed circle). Ribosomal RNA genes are transcribed at the FC-DFC interphase with the nascent rRNA extending into the DFC, which also harbors all rRNA modifying snoRNPs and Nopp140.

Knockdown of Nopp140 causes a specific loosening of the DFC (Bizarro et al., 2019, 2021). Appending GFP to PAF49 does not alter the tripartite structure of the nucleolus (Fig. 4I b; 1 and 2 refer to two independent clones). However, replacement or truncation of the PAF49 IDR with GFP disrupted the tripartite organization causing a loss of clear distinction between the three nucleolar compartments (Fig. 4I c and d). Quantification of the number of nucleolar sections with at least one FC-DFC unit demonstrated a significant difference between cells containing GFP-tagged full-length (b) and those with deleted (c) or partially truncated IDR (d) (Fig. 4H). Therefore, removal of the PAF49 IDR causes dissociation of the FC from the DFC, i.e., transcribing RNA polymerase I from the modifying snoRNPs.

We determined the impact of nucleolar disruption on rRNA 2’-O-methylation by performing RiboMethSeq (RMS) (Birkedal et al., 2015; Sharma et al., 2017; Marchand et al., 2016). Unexpectedly, RMS on our wild type HAP1 cells and those with GFP CRISPRed into the PAF49 locus to generate PAF49-GFP, PAF49ΔIDR-GFP, and PAF49Δ1/2IDR-GFP revealed a nearly unaltered 2’-O-methylation state of rRNA in those cells (Fig. 4J). Only 6 out of 107 methylation sites exhibited a more than 1.2-fold variation in degree of modification, up or down. Apparently, snoRNPs depend not solely on the PAF49 NLS-IDR (and Nopp140) for associating with transcribing RNA pol I to scan the nascent rRNA for modification sites.

### A second class of interacting IDRs in snoRNPs

Although removal of the NLS-IDR of PAF49, i.e., the CTD of pol I, disrupts the canonical tripartite structure of mammalian nucleoli, cell growth and rRNA modification proceed normally (Fig. 4). Therefore, snoRNPs must have means to concentrate around nascent rRNA in addition to the NLS-IDRs of NAP57, NOP56, and NOP58 mediated by Nopp140 (Fig. 1-3). Indeed, GAR1 of H/ACA RNPs and fibrillarin of C/D RNPs contain additional IDRs. These IDRs are rich in RGG repeats (Fig. 5A and B) that have been shown to interact with RNA and to phase separate (Girard et al., 1992; Ghisolfi et al., 1992; Kiledjian and Dreyfuss, 1992; Feric et al., 2016). Upon closer inspection, these IDRs are also enriched in phenylalanine (F) often preceded by glycine (G), i.e., in GF dipeptides, which sometimes expand to GFG and GFRG in various combinations (Fig. 5A and B). These GF repeats are reminiscent of the FG repeats of nucleoporins that form the permeability barrier of nuclear pore complexes between nucleus and cytoplasm (Rout et al., 2000). FG IDRs of nucleoporins interact with each other, alone or with the help of karyopherins that bind to FG repeats (Frey et al., 2006; Cowburn and Rout, 2023). Consequently, using our pulldown assay, we tested if the GF repeats of GAR1 and fibrillarin interacted with themselves and each other. Indeed, MBP-GAR1 pulled down GST-GAR1 (Fig. 5C, lane 4) in an IDR-dependent manner (lane 5). Similarly, MBP-fibrillarin pulled down GST-GAR1 (Fig. 5D, lane 4) in an IDR-dependent manner (lane 5). Even the GF-IDR of fibrillarin alone was sufficient for GST-GAR1 pulldown (lane 6). These data support the interaction of the GF-IDRs even in the absence of RNA. Surprisingly, MBP-GAR1 alone was sufficient for pulldown also of Nopp140 (Fig. 5E, lane 7) in a GF-IDR-dependent manner (lane 8) similar to MBP-NAP57 (lane 6). But the C-terminal IDR of GAR1 alone was insufficient to pulldown Nopp140 (Fig. 5E, lane 9). MBP-fibrillarin also pulled down Nopp140 (Fig. 5F, lane 6) in a GF-IDR-dependent manner (lane 7). Even the MBP-GF-IDR alone was able to interact with Nopp140 (lane 8) almost as well as MBP-NAP57 alone (lane 5). Surprisingly, the interaction of Nopp140 with those GF-IDRs occurred via its alternatingly charged repeat domain (Fig. 5G).

**Figure 5.**
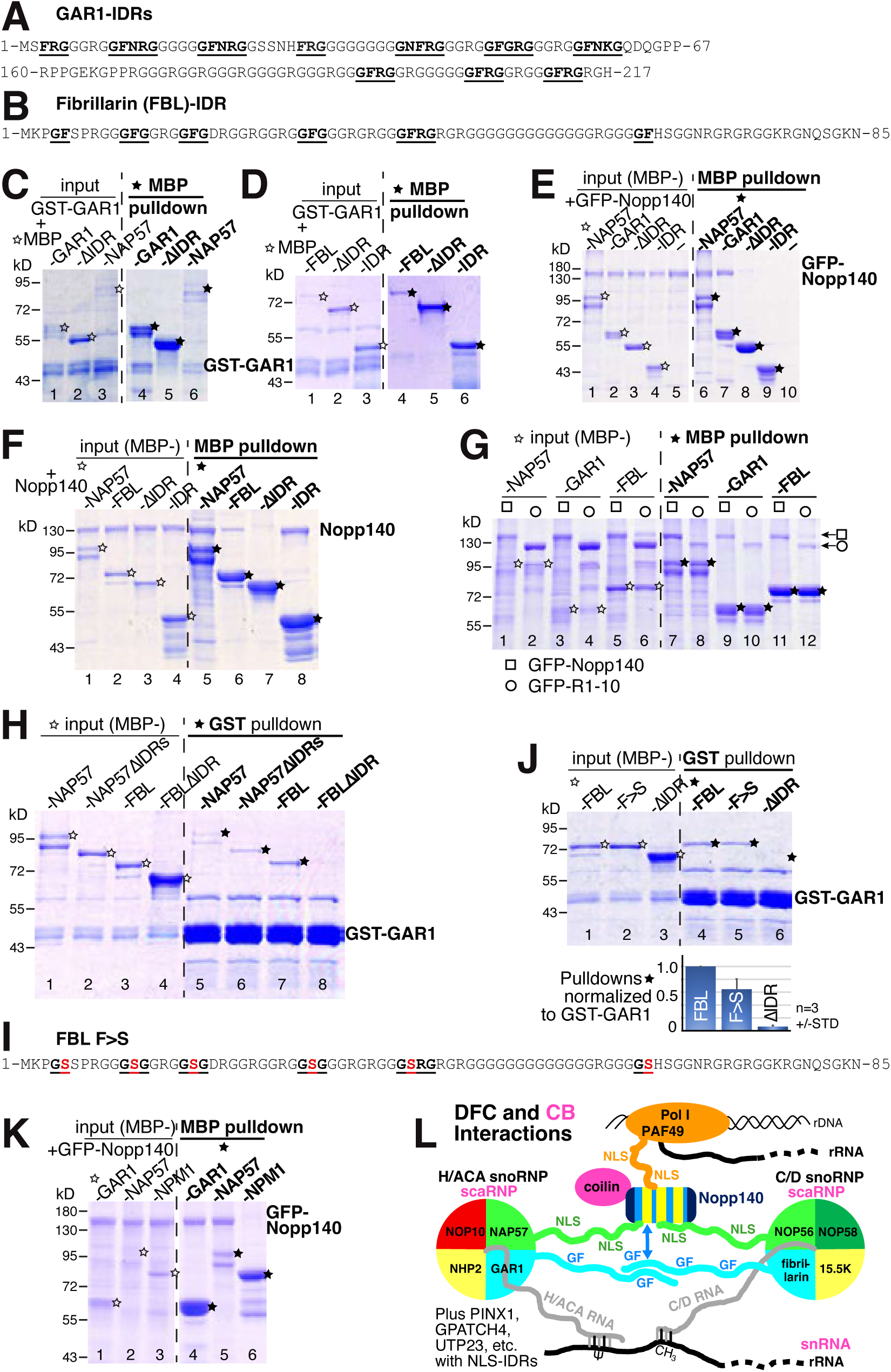
Interactions of GF repeat-IDRs. (A) Amino acid sequences of the N- and C-terminal glycine-arginine-rich IDRs of GAR1. GF repeats are highlighted (bold and underlined). (B) The N-terminal RGG IDR of fibrillarin with the GF repeats highlighted. (C) MBP pulldowns of GST-GAR1 with GAR1 (4), GAR1ΔIDR (5), and NAP57 (6) fused to MBP. All except GAR1ΔIDR pulldown GST-GAR1. (D) MBP pulldowns of GST-GAR1 with fibrillarin (FBL, 4), FBLΔIDR (5), and FBL-IDR alone (6) fused to MBP. All except FBLΔIDR pulldown GST-GAR1. (E) MBP pulldowns of GFP-Nopp140 with NAP57 (6), GAR1 (7), GAR1ΔIDR (8), and the GAR1-IDR alone (9) fused to MBP. Only full-length GAR1 (lane 7) but not without its IDR (lane 8) or the IDR alone pulled down GFP-Nopp140. (F) Similarly, MBP fused to NAP57 (lane 5), FBL (lane 6), and the FBL-IDR alone (lane 8), but not the FBL without its IDR (lane 7) pulldown Nopp140. (G) MBP pulldowns of GFP-Nopp140 (odd lanes, squares) and its repeats (even lanes, circles) with MBP-NAP57, -GAR1, and -FBL. (H) GST-GAR1 pulldowns of NAP57 and FBL with and without their IDRs. NAP57 (lane 6) but not FBL (lane 8) is pulled down by GST-GAR1 independent of its IDR. (J) The phenylalanines of the GF repeats of FBL are required for full interaction with GST-GAR1. Substitution of the phenylalanines by serines (F>S) reduces binding by 40% (lane 5) whereas IDR removal abolishes interaction. Quantification underneath. (I) Sequence of the FBL IDR with the serine substitutions (F>S). (K) Only DFC, but not GC, proteins interact with Nopp140. MBP pulldowns of GFP-Nopp140 with fusions of the DFC proteins GAR1 (lane 4) and NAP57 (lane 5) but not the GC protein nucleophosmin (NPM1, lane 6). (L) Schematic of the IDR interactions in the DFC of the nucleolus and in CBs as discussed in the text. The three major factors concentrating sno- or scaRNPs near their target RNAs are NLS-rich IDRs (via Nopp140), GF-rich IDRs (directly and via Nopp140), and the H/ACA and C/D RNA hybrids. In CBs (pink), PAF49 of pol I is replaced by coilin that directly interacts with Nopp140. The additional DFC proteins PINX1, GPATCH4, and UTP23 all contain C-terminal and NLS-rich IDRs.

MBP-GAR1 and MBP-fibrillarin pulled down the Nopp140 repeat domain alone (Fig. 5G, lanes 9-12) similar to MBP-NAP57 (lanes 7 and 8). To test the reverse interaction, we used GST-GAR1 to pull down NAP57 and fibrillarin constructs on glutathione beads (Fig. 5H). GAR1 pulled down both, NAP57 with and without its NLS-IDRs (Fig. 5H, lanes 5 and 6). This was to be expected as the H/ACA RNP structure shows that GAR1 and NAP57 to interact through their core domains independent of their IDRs (Ghanim et al., 2024). In contrast, GAR1 pulled down only full-length fibrillarin but not without its GF-IDR indicating interaction between their GF-IDRs (Fig. 5H, lanes 7 and 8).

Therefore, snoRNPs can directly interact with each other through their GF-IDRs (Fig. 5C, D, and H) and indirectly through Nopp140 (E-G). This is reminiscent of the central plug in nuclear pore complexes where FG repeats of nucleoporins interact with each other and with karyopherins that are also present in the plug (Frey et al., 2006; Cowburn and Rout, 2023). In nuclear pore complexes, FG repeats form a kind of hydrogel through π-π stacking interactions of the phenylalanines. This hydrogel and permeability barrier is broken when the phenylalanines of the repeats are substituted with serines (Frey et al., 2006). To test if the snoRNP GF-IDR interactions also depended on the phenylalanines, we replaced the six phenylalanines with serines in the fibrillarin N-terminal GF-IDR (Fig. 5I). Substitution of the 6 phenylalanines reduced fibrillarin binding to GST-GAR1 by 35% (Fig. 5J, lane 5, see graph below the gel), whereas removal of the GF-IDR abolished binding altogether (lane 6). Therefore, the GF-IDRs of snoRNP core proteins appear to function similarly to the FG repeats of nucleoporins.

All proteins and interactions assessed so far are confined to the DFC of the nucleolus. Hence, we tested another prominent nucleolar protein for interaction with Nopp140, nucleophosmin or NPM1. NPM1 is a protein of the granular component of the nucleolus. It forms a pentamer and about half of its protein sequence is intrinsically disordered. Unlike GAR1 and NAP57 (Fig. 5K, lanes 4 and 5), NPM1 fails to pull down Nopp140 (lane 6). Thus, our in vitro interactions reflect the nucleolar compartmentalization and support the physiological roles of these proteins.

## DISCUSSION

Ribosomal RNA modification is a major task of every cell. Mostly nascent rRNA is modified by some 100-200 small nucleolar ribonucleoproteins (snoRNPs), which each must scan the 13kb-long rRNA in its entirety not to miss a site for hybridization to guide modification. We uncover a dense yet flexible framework that underlies this process forming the DFC of the nucleolus. It consists of a multitude of weak transient interactions between snoRNPs, nascent rRNA, elongating RNA pol I, coalesced by the alternatingly charged chaperone Nopp140. Of course, additional ribosome assembly factors are present as well, some also interacting with Nopp140. Altogether, we describe the molecular interactions of a highly dynamic network that forms the liquid-liquid phase separated DFC. It mainly consists of intrinsically disordered regions (IDRs) of snoRNP core proteins and of PAF49, which are concentrated around nascent rRNA by Nopp140 with its own IDR.

There are three major forces holding the network together while permitting free mobility to snoRNPs and Nopp140 (Fig. 5L). First, the NLS-rich IDRs of both the H/ACA and C/D snoRNP core proteins NAP57, NOP56, and NOP58, and of the pol I CTD, PAF49, with all NLSs interacting with Nopp140. Since all IDRs have multiple NLSs and Nopp140 contains 10 alternatingly charged repeats, there appears to be an endless combination of interactions possible contributing to the overall concentration of snoRNPs. The complementary electrostatic interaction between the NLS IDRs and Nopp140 repeats is highly reminiscent of the spacers and stickers model underlying liquid-liquid phase separation (Das and Pappu, 2013; Brangwynne et al., 2009; Li et al., 2012; Martin et al., 2020; Harmon et al., 2017). These interactions may also provide the molecular explanation for an association between pol I and Nopp140 described some time ago (Chen et al., 1999). Second, the GF repeats of the GAR IDRs, which can interact with each other but also with Nopp140. The nature of the latter has yet to be determined, but two possibilities come to mind, the arginines of the GAR IDRs could interact with the negatively charged and phosphorylated serine stretches of the Nopp140 repeats or simply, IDRs in general exhibit affinity for each other (Martin and Holehouse, 2020). Third, snoRNA-rRNA hybrids mediate further interaction between snoRNPs and rRNA. This may also explain why, even in the absence of the PAF49 IDR, rRNA is nearly fully modified. Despite this multitude of interactions, they are all weak and transient allowing all components to move around rapidly forming a liquid-liquid biomolecular condensate that constitutes the DFC.

The specialized phase formed by the GF repeats and by the charged NLS IDRs with Nopp140 are highly reminiscent of nuclear transport and the central phase of the nuclear pore complex. In case of nuclear protein import, soluble nuclear transport receptors (NTRs) recognize the NLSs of their import cargo directly or indirectly. All NTRs of the beta karyopherin family can simultaneously bind to the FG repeats, IDRs of nucleoporins that face the central channel of the nuclear pore. This is similar to soluble Nopp140 binding to the NLSs of the charged IDRs of snoRNP and other DFC proteins. The central phase of the pore is made more permeable by mixing FG repeats with NTRs (Cowburn and Rout, 2023), though the FG repeats are anchored in the NPC through the nucleoporins that harbor them. Similarly, the DFC is defined by transcribing pol I molecules that are restricted to the rDNA genes at the FC-DFC interphase. Nopp140 then can mediate interaction between the PAF49 NLS-IDR and those of the snoRNP core proteins while the phase is further stabilized by NLS-IDRs of additional ribosome biogenesis factors, e.g., PINX1, GPATCH4, and UTP23 (Kanwal et al., 2023). Despite the anchoring of these IDRs in the NPC and at the elongating pol I, transiting molecules can readily slide through supporting these highly dynamic processes, i.e., transport across the NPC and scanning 13kb of pre-rRNA for cognate sites of snoRNAs.

Altogether, the IDRs of the proteins combined amass to some 1880 disordered amino acids per one H/ACA RNP, C/D RNP, Nopp140, PAF49, PINX1, GPATCH4, and UTP23 in the DFC alone. Of course, the stoichiometry will differ greatly as Nopp140 with 10 repeats and PAF49 with 9 NLSs alone can accommodate multiples of each other. Removing the GF repeats of GAR1 and fibrillarin and the Nopp140 repeats still leaves us with 1170 disordered amino acids of charged IDRs that harbor 37 NLSs. We show that Nopp140 is responsible for concentrating the close to 200 snoRNPs around elongating pol I via PAF49. Surprisingly though, the PAF49 IDR is not essential for rRNA modification in the nucleolus, nor in CBs, which lack PAF49, yet mediate snRNA modification. The small but detectable difference in modification with and without the PAF49 IDR may reflect the small difference observed when transcription of the target RNA is driven by pol I versus pol II (Nir et al., 2022). Although PAF49 is not an integral part of pol I, it exhibits remarkable similarities to the pol II integral C-terminal domain (CTD). The latter consists of 52 seven-amino acid repeats amounting to 364 residues whereas the PAF49 CTD harbors 9 NLS repeats separated by negatively charged spacers adding up to 350 residues, i.e., similar in length. Both are intrinsically disordered and function as landing platforms for factors required for integrating transcription with RNA processing and modification (Brickey and Greenleaf, 1995; Meinhart and Cramer, 2004; Corden, 1990). Although only loosely related on an evolutionary scale, these IDRs are similarly important for nucleolar compaction in yeast. Removal of the charged IDRs of Nop56 and Nop58 in yeast causes a loosening of the nucleolus (Colau et al., 2004; Dominique et al., 2024) like Nopp140 knockdown in mammalian cells (Bizarro et al., 2021, 2019).

CBs are closely related to DFCs, which are the only nuclear bodies in the cell to concentrate both Nopp140 and sno- or scaRNPs (Fig. 5L). The main function of CBs is to serve as location for snRNA modification by scaRNPs (Bizarro et al., 2021). Apparently, scaRNPs are concentrated in CBs by interaction with Nopp140 like snoRNPs in the DFC. Nopp140 additionally binds to coilin the CB scaffold protein (Isaac et al., 1998; Courchaine et al., 2022). Upon Nopp140 knockdown, scaRNPs are specifically lost from CBs and disperse into the nucleoplasm causing a loss of most snRNA modification (Bizarro et al., 2021). The loss of scaRNPs is supported by a shrinking by half of the granules making up CBs, which is similar to the loosening of the DFC after Nopp140 knockdown (Bizarro et al., 2019, 2021). Apparently, Nopp140 concentration of sno- and scaRNPs contributes equally to liquid-liquid phase separation and biomolecular condensate formation of DFCs and CBs, respectively. DFCs are tethered to rDNA genes in the nucleus, but CBs are freely floating in the nucleoplasm though sometimes appear near snRNA gene clusters (Frey and Matera, 1995; Smith et al., 1995).

One of the most unexpected results of our study was the lack of effect of removal of the C-terminal IDR of PAF49 despite causing the disruption of nucleolar organization (Fig. 4I and H). The strict organization of the DFC surrounding the FC is lost. However, individual pieces, but not units of FCs and DFCs can be recognized. This may explain why these cells divide at equal rates and why rRNA modification is barely affected. Thus, snoRNPs are not only concentrated around nascent rRNA by binding to Nopp140 and the PAF49IDR, but also through snoRNA-rRNA base pairing as depicted (Fig. 5L). The snoRNP IDRs in general aid the concentration of the snoRNPs directly or indirectly through Nopp140. Apparently, PAF49 is mostly required for the stabilization of its essential heterodimeric partner PAF53, which occurs in the absence of its IDR (Penrod et al., 2012; Yamamoto et al., 2004; McNamar et al., 2023). Even rRNA 2’-O-methylation is barely affected under those conditions (Fig. 4J). Although the nucleolar tripartite structure is conserved in mammalian cells, it is not essential for ribosome synthesis and cell growth.

Overall, we describe in molecular detail the network that underlies the DFC, a highly dynamic biomolecular condensate. The crucial role of Nopp140 and IDRs in this process is supported by Nopp140 knockdown and IDR removal and mutation.

## MATERIALS AND METHODS

### Plasmids

All plasmids used in this study are described in table S1. Sequences encoding human nucleolar proteins used in this study were either transferred by direct cloning from existing plasmids or amplified by PCR before insertion into the corresponding vectors (see next paragraph). Plasmids were sequenced after cloning. Primers used for PCR amplification and sequencing are listed in table S2.

The following plasmids served as templates for CDS amplification: for PAF49, pBAD-CAST from Kostya Panov (Queens University, Belfast); for PINX1, pRC43 from Yves Henry (University of Toulouse, Toulouse, Fance); for nucleophosmin, pRUTH_NPM1 from David Shechter (Albert Einstein College of Medicine, Bronx, USA); for Nop56 and Nop58, peGFP-hNop56FL and peGFP-hNop58FL, respectively, from Edouard Bertrand (Institute of Human Genetics, Montpellier, France), for GAR1, pAD-hGAR1 from Francois Dragon (Université du Québec à Montréal, QC, Canada); and GFP-Fibrillarin (Platani et al., 2000).

The construction of each Knock-In plasmid (used for genome engineering) consisted of two successive cloning steps. First, annealed oligos encoding a guide RNA were inserted into pORANGE (Willems et al., 2020). Second, monomeric GFP amplified by PCR with primers fused to an inverted target sequence of each corresponding gRNA. Primers used for the cloning are listed in table S2.

### Protein purification

Except for Nopp140 derived proteins, which were expressed in SF9 cells, all recombinant proteins were expressed in bacterial strain Rosetta 2 (Novagen). Using the vectors listed in table S1, Nopp140 or fragments thereof were expressed in SF9 cells using Bac-to-Bac^TM^ Baculovirus Expression System (Gibco) following the protocol of the supplier. After collection of Nopp140-expressing SF9 cells, Nopp140 fragments (all tagged with 6xHis) were purified by affinity chromatography on Nickel-NTA resin. Briefly, cell pellets were resuspended of 1/20^th^ culture volume in lysis buffer (50 mM Tris-pH7.9, 100 mM KCl, 0.5% (w/v) IGEPAL CA-630, 0.2 mM EDTA, 10 mM Imidazole, 1x EDTA-free Pierce Protease Inhibitor cocktail), sonicated 4x times for 20 seconds, the lysate cleared by sedimentation at 34,000 x *g* for 20 minutes at 4°C, and loaded onto a Nickel-NTA resin column. After washing, the proteins were eluted with 20 mM Tris-pH7.9, 150mM NaCl, 250 mM Imidazole and dialyzed against 20 mM Tris-pH7.9, 150mM NaCl.

Transformed Rosetta 2 strains were grown at 37°C and expression induced with 1 mM IPTG either for 3 h at 37°C (MBP-MCP, MBP-NMP1-His, MBP-FibrillarinRGG-His, MBP-GAR1-His, MBP-GAR1 Core-His, MBP-PAF49-His, MBP-PAF49 ΔIDR-His and MBP-PAF49 IDR-His proteins) or overnight at 16°C (all other proteins). Cells were resuspended in 1/20th culture volume of 50 mM Tris-pH7.4, 500 mM NaCl, 10mM imidazole in case of nickel NTA purifications, (1mM EDTA in case of amylose chromatography), supplemented with 1x EDTA-free Pierce Protease Inhibitor cocktail then lysed by nitrogen cavitation through an EmulsiFlex-C5 (Avestin Inc.). Proteins were affinity purified using Nickel-NTA resin (Qiagen), amylose resin (New England Biolabs) or glutathione sepharose beads (Pierce^TM^) as listed in table S1. For dual affinity purified proteins like MBP-NAP57-His, purifications were carried out as previously described in (Grozdanov et al., 2009) but with amylose resin chromatography being performed last. All proteins purified from bacteria were dialyzed against 20 mM Tris-pH7.4, 300mM NaCl.

### In vitro pulldown assays

For pulldowns, bait proteins (MBP- or GST-tagged proteins) and their partners were incubated for 30 min at room temperature in the presence of 150 mM (or 120 mM in Figs 1B-D and 1F-G) NaCl. The reactions were then added to 25 µL of the corresponding resin, i.e. amylose (New England Biolabs) or glutathione (Pierce) in ∼200 µL binding buffer (150 mM NaCl; 20 mM Tris-HCl, pH 7.4; 0.02% TritonX-100). After 30 min incubation at room temperature, the resins were washed 3 times with 1 ml of binding buffer, resuspended in Laemmli buffer then visualized by coomassie blue staining after SDS-PAGE. Fractions corresponding to 5% of the initial input were also analyzed. Quantifications were performed using Image Studio Lite (LI-COR Biosciences) and normalized as ratios of proteins over bait proteins. Most pulldowns were performed at least three times with identical results.

### Cell culture, transfection and genome engineering

U2OS cells with integrated LacO arrays (Janicki et al., 2004) were cultured in DMEM (Corning), while HAP1 cells (Carette et al., 2011) and its derived clones were cultured in HyClone^TM^ Iscoves’s Modified Dulbecco’s Medium (IMDM, Cytiva). Both media were supplemented with 10% fetal bovine serum (Atlanta Biologicals) and all cells cultured at 37°C under 5% CO2 in air.

Plasmid transfections were carried out using Lipofectamine 3000 (InVitrogen) following the manufacturer’s protocol. For in vivo tethering assays, U2OS cells were transfected for 24 hours with the vectors expressing the Lac-repressor (LacI) fused to monomeric RFP with or without the different IDRs (see vector details in table S1).

Genome engineering strategy to delete parts or all of the PAF49 IDR consisted of insertion of monomeric GFP while replacing the IDR. Thus, insertion of mGFP at 3 sites yielded cells expressing PAF49 full length-mGFP, PAF49 ΔIDR-mGFP or PAF49 ½ IDR-mGFP. Genome engineering was carried out using the CRISPR/Cas9 system and mGFP integration was achieved using a homology-independent targeted integration (HITI) strategy (Suzuki et al., 2016). The pORANGE vector (Willems et al., 2020) was used to generate knock-in plasmids containing all the required elements for the HITI strategy, i.e., expression of the Cas9 protein and PAF49-targeting guide RNAs while also carrying the donor sequence (mGFP) enclosed between two inverted target sequences, i.e., recognized by the guide RNA. The sequences of guide RNAs of the three different knock-in vectors are listed in table S1.

Each knock-in plasmid was co-transfected with an mRFP-C1 vector in a ratio of 2:1 for 2 days before FACS sorting of RFP-positive cells. Sorted cells were cultured for 7-8 days before single cell sorting of GFP+, i.e., knock-in cells. After cell expansion, knock-in clones were checked for proper insertion by PCR and sequencing using the primers listed in table S2. For growth evaluation, 12,000 cells of each clone were plated on day 1 and expansion assessed on 4 subsequent days using a Countess II automated cell counter (Thermo Fisher Scientific).

### Antibodies

The following antibodies were used for Western blotting (WB) or indirect immunofluorescence (IF): anti-Nopp140 rabbit serum (RS8 at 1:5000 for WB; 1:1000 for IF) (Kittur et al., 2007); mouse monoclonal anti-NAP57 immunoglobulin G (IgG) (H3 at 1:500 for IF; Santa Cruz Biotechnology); anti-NAP57 rabbit serum (RU10 at 1:200 for WB) (Darzacq et al., 2006); mouse monoclonal anti-fibrillarin culture supernatant (38F3 at 1:2000 for WB; EnCor Biotechnology); mouse monoclonal anti-PAF49 (MA5-27813 at 1:1000 for WB; 1:500 for IF; Invitrogen); rabbit polyclonal anti-GFP (ab290 at 1:1000 for WB; Abcam); DyLight488 goat anti-mouse IgG and rhodamine (TRITC) goat anti-rabbit IgG (1:500 for IF; both Jackson Immuno Research); Alexa Fluor 647 goat anti-mouse IgG (1:500 for IF) and Alexa Fluor 680 goat anti-rabbit IgG (1:10,000 for WB; from Thermo Fisher Scientific); IRDyeTM 800 goat anti-mouse IgG (1:10,000 for WB; Rockland Immunochemicals).

### Immunofluorescence

For immunofluorescence, cells were grown on coverslips and cultured for 1 day before being transfected (U2OS cells) or directly fixed. Cells were washed with PBS then fixed with 4% paraformaldehyde for 20 min at room temperature. After being washed twice with PBS, cells were permeabilized with 1% (w/v) Triton-X100 in PBS for 5 min before 3 washes. Cells were blocked for 15 min at room temperature with blocking buffer (PBS, 1% nonfat powdered milk), then probed with primary antibodies (diluted in blocking buffer) for 2h. Following three washes with blocking buffer, cells were probed with secondary antibodies for 1h in the dark, washed twice, counterstained for 5 min with 4′,6-diamidino-2-phenylindole (DAPI), washed with PBS, and the coverslips mounted on glass slides using ProLong Diamond Antifade Mount (Thermo Fisher Scientific). Image acquisition and analysis were performed as described (Bizarro et al., 2021).

### Western blotting

Cells were lysed in 1X Laemmli buffer and sonicated 10 seconds for 5 times. Total protein extracts, equivalent to 60,000 cells, were separated by SDS-PAGE and transferred to a nitrocellulose membrane at 100 volts for 1h. Membranes were blocked with 2.5% (w/v) powdered milk in PBS for 40 min at room temperature and probed with primary antibodies prepared in dilution buffer (PBS, 0.1% Tween, 1% nonfat dry milk) for 2h at room temperature or overnight at 4°C. After three washes in dilution buffer, membranes were incubated with appropriate secondary antibodies diluted in blocking buffer for 1 h at room temperature in the dark. Membranes were washed three times in blocking buffer, scanned on an Odyssey CLx imaging system model 9120 (LI-COR Biosciences), and protein bands were quantified using Image Studio Lite (LI-COR Biosciences).

### Electron microscopy

Transmission electron microscopy was performed exactly as described (Bizarro et al., 2019). Briefly, cells were fixed with 2.5% glutaraldehyde in 0.1 M sodium cacodylate buffer, postfixed with 1% osmium tetroxide followed by 2% uranyl acetate, dehydrated through a graded series of ethanol, and embedded as a loose pellet in LX112 resin (LADD Research Industries, Burlington, VT). Ultrathin sections were cut on a Leica Ultracut UC7, stained with uranyl acetate followed by lead citrate, and viewed on a JEOL 1400 Plus transmission electron microscope at 80 kv.

### Analysis of 2′-O-methylation levels by RiboMethSeq

RMS was performed exactly as described (Marchand et al., 2016). For each reaction, 150 ng of total RNA was used. The samples were sequenced at the ULB-BRIGHTcore facility (Brussels Interuniversity Genomics High-Throughput Core) on Illumina Novaseq 6000 as paired-end runs (100-nt read length). On average, 25 million reads were sequenced. Adapter sequences were removed using Trimmomatic (0.36; leading:30 trailing:30 slidingwindow:4:15 minlen:17 avgqual:30) and reads in forward direction were mapped to an artificial genome containing ribosomal RNA sequences using bowtie2 (2.3.3.1; sensitive). Mapped reads were analyzed using the R package version 1.2.0 RNAmodR. RiboMethSeq (https://bioconductor.org/packages/release/bioc/html/RNAmodR.RiboMethSeq.html) and the Score C/RiboMethScore was used as a measurement for 2′-O methylation. For the analysis of methylation levels on positions known to be methylated, data from the snoRNAdb (Lestrade and Weber, 2006) were used and updated to revised rRNA sequence coordinates based on NCBI accession NR_046235.3, which are available from the EpiTxDb R package version 1.0.0 (http://www.bioconductor.org/packages/release/bioc/html/EpiTxDb.html).

## ACKNOWLEDGEMENTS

We are grateful to all people mentioned in the methods section for providing plasmids and antibodies.

