## Supplemental Table 1 and 2 for "Intrinsically Disordered Regions Form Nucleoli and Cajal Bodies While Fostering RNA Modification"

TABLE S1

List of vectors
(all constructs are based on human protein sequences)

Table S1a. Vectors for recombinant protein purification

| | **Vector** | **Protein expressed** | **Description** | **Affinity purification resin** |
|---|---|---|---|---|
| for protein expression in bacteria | pMAL-MCP | MBP-MCP | Grozdanov et al, 2009a | Amylose |
| | pPG17 | MBP-NAP57-His | Grozdanov et al, 2009a, expression of NAP57 FL (amino acid 1-514) | Nickel-NTA + Amylose |
| | pTM193 | MBP-NAP57 ΔN & ΔC -His | Walbott et al, 2011, Gene & Dev, (NAP57 aa 31-422) | Nickel-NTA + Amylose |
| | pRM67 | MBP-NAP57 ΔC-His | Machado-Pinilla et al, 2012, RNA, NAP57 (aa 1-422) | Nickel-NTA |
| | pRM68 | MBP-NAP57 ΔN-His | Machado-Pinilla et al, 2012, RNA, (NAP57 aa 31-514) | Nickel-NTA |
| | pMD26 | MBP-NAP57 C-His | NAP57 C (aa 423-514) amplified from pPG17 using U856/U857, and inserted in the same backbone as pPG17 | Nickel-NTA |
| | pMD35 | MBP-NAP57 N-His | NAP57 N (aa 1-30) amplified from pPG17 using U392/U896, and inserted in the same backbone as pPG17 | Amylose |
| | pMD17 | MBP-Nop56-His | Nop56 CDS amplified from peGFP-hNop56FL (provided by Edouard Bertrand) using U811/U812 and inserted in the same backbone as pPG17 | Nickel-NTA then Amylose |
| | pMD22 | MBP-Nop56 ΔC-His | Nop56 ΔC (aa 1-433) CDS amplified from peGFP-hNop56FL (provided by Edouard Bertrand) using U811/U826 and inserted in the same backbone as pPG17 | Nickel-NTA |
| | pMD28 | MBP-Nop56 C-His | Nop56 C (aa 434-514) CDS amplified from peGFP-hNop56FL (provided by Edouard Bertrand) using U858/U812 and inserted in the same backbone as pPG17 | Nickel-NTA |
| | pMD18 | MBP-Nop58-His | Nop58 CDS amplified from peGFP-hNop58 FL (provided by Edouard Bertrand) using U813/U814 and inserted in the same backbone as pPG17 | Nickel-NTA |
| | pKM171 | MBP-Nop58 mut all NLS-His | Replace Nop58 IDR IDR CDS from pKM109 by Lysine>Alanine mutants in all NLSs (Fig. 3C), synthetized (Twist Bioscience) | Nickel-NTA |

| | | | | |
|---|---|---|---|---|
| | pKM172 | MBP-Nop58 1 NLS-His | Replace Nop58 IDR CDS from pKM109 by Lysine>Alanine mutants in all but 1 NLSs (Fig. 3C), synthetized (Twist Bioscience) | Nickel-NTA |
| | pKM92 | MBP-PAF49-His | PAF49 CDS amplified from pBAD-CAST (provided by Joost CBM Zomerdijk) using U945/U946 and inserted in the same backbone as pPG17 | Nickel-NTA |
| | pKM93 | MBP-PAF49 ΔC-His | PAF49 ΔC amplified from pBAD-CAST (provided by Joost CBM Zomerdijk) using U945/U947 and inserted in the same backbone as pPG17 | Nickel-NTA |
| | pKM120 | MBP-PAF49 C-His | PAF49 C amplified from pBAD-CAST (provided by Joost CBM Zomerdijk) using U966/U947 and inserted in the same backbone as pPG17 | Nickel-NTA |
| | pRC43 | GST-PINX1-His | Provided by Yves Henry | Nickel-NTA |
| | pH177 | GST-PINX1 cat mut-His | Provided by Yves Henry | Nickel-NTA |
| | pKM112 | GST-PINX1 ΔC-His | Deletion of PINX1 IDR CDS from pRC43 by mutagensis using U961/U962 | Nickel-NTA |
| | pKM113 | Nop56-His | Deletion of MBP CDS from pMD17 by mutagensis using U963/U964 | Nickel-NTA |
| | pKM148 | MBP-NPM1-His | NPM1 CDS amplified from pRUTH_NPM1 (provided by David Shechter) using U992/U993 and inserted in the same backbone as pPG17. Nucleophosmin expression | Nickel-NTA |
| | pKM122 | GST-PAF49-His | Transfer of PAF49-His CDS from pKM92 into pGEX-KG | Nickel-NTA |
| | pKM89 | MBP-GAR1-His | GAR1 CDS amplified from pADhGAR1 (From Francois Dragon) using U941/U942 and inserted in the same backbone as pPG17 | Amylose |
| | pKM90 | MBP-GAR1 Core-His | GAR1 Core(ΔIDR) CDS amplified from pADhGAR1 (From Francois Dragon) using U943/U944 and inserted in the same backbone as pPG17 | Nickel-NTA |
| | pKM109 | MBP-Fibrillarin-His | Fibrillarin CDS amplified from GFP-Fibrillarin (Platani et al. 2000) using U958/U959 and inserted in the same backbone as pPG17 | Nickel-NTA |
| | pKM110 | MBP-Fibrillarin ΔRGG-His | Fibrillarin ΔGAR CDS amplified from GFP-Fibrillarin (Platani et al. 2000) using U960/U959 and inserted in the same backbone as Ppg17 | Nickel-NTA |
| | pKM156 | MBP-FBL RGG-His | Fibrillarin GAR CDS amplified from GFP-Fibrillarin (Platani et al. 2000) using U958/U1001 and inserted in the same backbone as pPG17 | Amylose |

| | | | | |
|---|---|---|---|---|
| | pKM121 | GST-GAR1-His | Transfer of GAR1-His CDS from pKM89 into pGEX-KG | Glutathione |
| | pKM180 | MBP-Fibrillarin FS-His | Replace RGG domain CDS from pKM109 by Phenylalanine>Serine mutant RGG (Fig. 5I) synthetized by TWIST Bioscience | Nickel-NTA |
| for protein expression in SF9 cells | pMD01 | His-Nopp140 | Insert Nopp140 coding sequence (KIAA0035) into pFastBacHT-b (Gibco) | Nickel-NTA |
| | pMD24 | His-GFP-Nopp140 | Insert superfolder GFP CDS amplified from pHR-SCFV-GCN4-sfGFP-GB1-NLS-dW (provided by Robert H Singer Lab) using U773/U879 into pMD01 | Nickel-NTA |
| | pKM86 | His-Nopp140 SA | Nopp140 mutant CDS (Serine>Alanine in Repeat domain) synthetized from Twist Bioscience and inserted in pFastBacHT-b | Nickel-NTA |
| | pKM88 | His-NoppR (R1-10) | Insertion of Nopp140 repeat domain (aa 65-581) into pFastBacHT-b | Nickel-NTA |
| | pKM98 | His-GFP-NoppR (R1-10) | Insertion of supefolder GFP CDS amplified from pMD24 using U773/U879 into pKM88 | Nickel-NTA |
| | pKM104 | His-GFP-NoppR R1-7 | Deletion of Nopp140 CDS Repeat 8-10 from pKM98 by mutagensis using U952/U953 | Nickel-NTA |
| | pKM105 | His-GFP-NoppR R1-3 | Deletion of Nopp140 CDS Repeat 4-10 from pKM98 by mutagensis using U954/U955 | Nickel-NTA |
| | pKM128 | His-GFP-NoppR R1 | Deletion of Nopp140 CDS Repeat 2-10 from pKM98 by mutagensis using U952/U971 | Nickel-NTA |
| | pKM137 | His-GFP-NoppR R8-10 | Deletion of Nopp140 CDS Repeat 1-7 from pKM98 by mutagensis using U976/U978 | Nickel-NTA |

Table S1b. Vectors used for in vivo tethering assay.

| | **Vector** | **Protein expressed** | **Description** |
|---|---|---|---|
| for *in vivo* tethering assay | pKM135 | LacI-mRFP | Lac repressor CDS amplified from pJRC70 (described in Darzacq et al. 2006, a gift from J.Chubb, University of Dundee, UK) using U973/U974, and inserted into mRFP-C1 |
| | pKM147 | LacI-mRFP-PAF49 IDR | PAF49 IDR amplified from pKM92 using U966/U948, and inserted into pKM135 |
| | pKM194 | LacI-mRFP-Nop58 IDR | Nop58 IDR amplified from pMD18 using U1019/U1020, and inserted into pKM135 |
| | pKM195 | LacI-mRFP-Nop58 IDR mut all NLS | Nop58 mut all NLS amplified from pKM171 using U1021/U1020, and inserted into pKM135 |

Table S1c. Vectors used for genome engineering.

| genome engineering vectors | Vector | Expressed protein and gRNA (annealed oligos inserted) | Donor sequence | Knock-In product |
|---|---|---|---|---|
| | pKM144 | Cas9 and gRNA KM1 (Annealed U986/U987) | mGFP flanked with inverted gRNA KM1 sites, amplified from mGFP-C1 using U980/U981 primers | PAF49 ΔIDR-GFP |
| | pKM146 | Cas9 and gRNA KM26 (Annealed U990/U991) | mGFP flanked with inverted gRNA KM26 sites, amplified from mGFP-C1 using U984/U985 primers | PAF49 1/2IDR-GFP |
| | pKM152 | Cas9 and gRNA 8 (Annealed U994/U995) | mGFP flanked with inverted gRNA 8 sites, amplified from mGFP-C1 using U996/U997 primers | PAF49-GFP |

TABLE S2

Oligonucleotides

| Oligo | Sequence | Description | Application |
|---|---|---|---|
| U986 | caccgGACTGGTGGGTTGCCCCCAA | Annealead then inserted into pORANGE, allowing for sgRNA KM1 expression | Construction of Knock-In plasmid pKM144 |
| U987 | aaacTTGGGGGCAACCCACCAGTCc | | |
| U980 | atgcAAGCTTgactggtgggttgcccccaaaggATATGGTGAGCAAGGGC | Amplification of mGFP flanked with gRNA KM1 target sequence with HindIII and XhoI sites | |
| U981 | atgcggatcccctttgggggcaacccaccagtcTTACTTGTACAGCTCGTCC | | |
| U990 | caccgTCTTCTTCGTAGGGGGCAGA | Annealead then inserted into pORANGE, thus allowing for sgRNA KM26 expression | Construction of Knock-In plasmid pKM146 |
| U991 | aaacTCTGCCCCCTACGAAGAAGAc | | |
| U984 | atgcAAGCTTTCTTCTTCGTAGGGGGCAGAGGGAATGGTGAGCAAGGGC | Amplify of mGFP flanked with gRNA KM26 target sequence with HindIII and XhoI sites | |
| U985 | atgcGGATCCccctctgcccccctacgaagaagaTTACTTGTACAGCTCGTCC | | |
| U994 | caccgttcccgggggcagaCTACAC | Annealead then inserted into pORANGE, thus allowing for sgRNA 8 expression | Construction of Knock-In plasmid pKM152 |
| U995 | aaacGTGTAGtctgcccccgggaac | | |
| U996 | atgcAAGCTTttcccgggggcagaCTACACAGGATGGTGAGCAAGGGC | Amplify of mGFP flanked with gRNA 8 target sequence with HindIII and XhoI sites | |
| U997 | atgcggatccCCTGTGTAGtctgcccccgggaaTTACTTGTACAGCTCGTCC | | |
| U1002 | CTGTCCCCAAGCTGGAGAAG | PCR screening of PAF49ΔIDR-GFP cells | |
| U975 | agtcaCTCGAGttctagaggctcagtgttaat | | |
| U988 | caccgtCCCGTCCACCACCAAGAAG | PCR screening of PAF49-GFP and PAF491/2IDR-GFP cells | |
| U998 | TGGGCACCTTCAGCTTTCTT | | |
| GFP-F | CCGACAACCACTACCTGAGC | Sequencing of integration sites (3' junction) | |
| GFP-R | GCTCGTCCATGCCGAGAGTG | Sequencing of integration sites (3' junction) | |
| U856 | agggcagatctgccaaaaaagaggtggt | Amplification of NAP57 C CDS | |
| U857 | taagacctcgagctcagaaaccaattctacct | | |
| U392 | AGGGCagatctATGGCGGATGCGGAAGTAATT | Amplification of NAP57 N CDS | |
| U896 | ctcgagTCAtatttcggctacatcttctt | | |
| U811 | agggcagatctATGGTGCTGTTGCACGTGCTGTTTG | Amplification of Nop56 | |
| U812 | taagacctcgaggccATCTTCCTGGGATGCTTTATGG | Amplification of Nop56 | |

| | | |
|---|---|---|
| U826 | taagacctcgaggccAGCAGCCGCTTC | Amplification of Nop56 ΔC |
| U858 | agggcagatctGAGATTACTAGGAAGCT | Amplification of Nop56 C |
| U813 | CAGGGCggatccATGTTGGTGCTGTTTGAAACGTCTG | Amplification of Nop58 CDS |
| U814 | taagacctcgaggccATCCTCGTTCTCTCTC | |
| U945 | gatcAGATCTatggaggagccccag | Amplification of RPA34 |
| U946 | taccCTCGAGcacaggctgctgc | Amplification of RPA34 |
| U947 | agtcCTCGAGccctgaggtgagcagg | Amplification of RPA34 ΔC |
| U966 | agctcAGATCTgggaagaagaaaaaggagatg | Amplification of RPA34 C |
| U961 | CACCATCACCATCACCATTAAGC | Deletion of PINX1 C-terminus |
| U962 | CAGTGCTGCCATCCGCTT | |
| U963 | TTTCAGGGCGGATCCATG | Deletion of MBP CDS |
| U964 | CATAATCTATGGTCCTTGTTGG | |
| U992 | atgcGGATCCatgGAAGATTCGATGGACATGGAC | Amplification of NPM1 CDS |
| U993 | agtcctcgagAAGAGACTTCCTCCACTGC | |
| U941 | gatcGGATCCatgtcttttcgaggcggag | Amplification of GAR1 CDS |
| U942 | gatcCTCGAGatgtcctctccctctgaaac | |
| U943 | gatcGGATCCATGcaagaccaaggacctccag | Amplification of GAR1 Core CDS |
| U944 | gatcCTCGAGtggaggtcctttctcacc | |
| U958 | aatctGGATCCatgaagccaggattcagtcc | Amplification of Fibrillarin |
| U959 | aatctCTCGAGgttcttcaccttgggggg | Amplification of Fibrillarin |
| U960 | aatctGGATCCgtgatggtggagccgc | Amplification of Fibrillarin ΔRGG |
| U1001 | aatctCTCGAGatgcggctccaccatcac | Amplification of Fibrillarin RGG |
| U773 | ggcgccATGGCGAGCAAAGGAGAAGAAC | Amplification of sfGFP |
| U879 | TATACCATGgTGCCCTGAAAATACAGGTTTTCTTTGTAGAGCTCATCCATGCCA | |
| U748 | GATACAGGTCGCTGGGAACC | |
| U952 | AAGGCGGCAGTGGTAGTT | Deletion of Nopp140 Repeat 8-10 or 2-10 |
| U953 | AGAATTCTTGGTGGTACCAGC | Deletion of Nopp140 Repeat 8-10 |
| U971 | TGATGCTTTGGCTGCAGC | Deletion of Nopp140 Repeat 2-10 |
| U954 | aagAATTCTaaggcggcagtggtagtt | Deletion of Nopp140 Repeat 4-10 |
| U955 | ggtggtaccagcaggtgctgctcgagc | |
| U976 | TCAAATAAGCCAGCTGTC | Deletion of Nopp140 Repeat 1-7 |
| U978 | TCCATTTGCCTGTAACTTTC | |
| U973 | atcaccggtgccaccatgggcAAACCAGTAACGTTATACG | Amplification of LacI |
| U974 | atgcaccggtccTCGGGAAACCTGTCGTGCCA | |
| U948 | TATTAaagcttTCAcacaggctgctgctg | Amplification of PAF49 IDR |

| | | |
|---|---|---|
| U1019 | atgcagatctTCTAAAAAACGCAAAATAGAA CAG | Amplification of Nop58 IDR |
| U1020 | atgcaagctttcaATCCTCGTTCTCTCTCTT | Amplification of Nop58 IDR WT or mutant |
| U1021 | atgcagatctTCTAAAgcACGCAAAATAGAA CA | Amplification of Nop58 mut IDR |
